# A gas phase fractionation acquisition scheme integrating ion mobility for rapid diaPASEF library generation

**DOI:** 10.1101/2022.07.21.500948

**Authors:** Jack Penny, Gunnar N. Schroeder, José A. Bengoechea, Ben C. Collins

## Abstract

Data independent acquisition (DIA or DIA/SWATH) mass spectrometry has emerged as a primary measurement strategy in the field of quantitative proteomics. diaPASEF is a recent adaptation that leverages trapped ion mobility spectrometry (TIMS) to improve selectivity and increase sensitivity. The complex fragmentation spectra generated by co-isolation of peptides in DIA mode are most typically analyzed with reference to prior knowledge in the form of spectral libraries. The best-established method for generating libraries uses data dependent acquisition (DDA) mode, or DIA mode if appropriately deconvoluted, often including offline fractionation to increase depth of coverage. More recently strategies for spectral library generation based on gas phase fractionation (GPF), where a representative sample is injected serially using narrow DIA windows that cover different mass ranges of the complete precursor space, have been introduced that performed comparably to deep offline fractionation-based libraries for DIA data analysis. Here, we investigated whether an analogous GPF-based library building approach that accounts for the ion mobility (IM) dimension is useful for the analysis of diaPASEF data and can remove the need for offline fractionation. To enable a rapid library development approach for diaPASEF we designed a GPF acquisition scheme covering the majority of multiply charged precursors in the m/z vs 1/K0 space requiring 7 injections of a representative sample and compared this with libraries generated by direct deconvolution-based analysis of diaPASEF data or by deep offline fractionation and ddaPASEF. We found that library generation by IM-GPF outperformed direct deconvolution of the diaPASEF data and had performance approaching that of a deep offline fractionation library, when analysing diaPASEF data. This establishes the ion mobility integrated GPF scheme as a pragmatic approach to rapid library generation for the analysis of diaPASEF data.

## Introduction

Data Independent Acquisition (DIA or DIA/SWATH) methods are becoming increasingly popular in quantitative proteomics studies because they provide a robust and flexible method for the generation of large-scale quantitative proteomics data [1–4]. DIA methods achieve this by deterministically acquiring MS2 spectra from co-fragmenting peptides within wide predefined precursor isolation windows that collectively span the m/z range of interest [5]. Typical computational workflows for analysis of DIA data have required the generation of reference spectral libraries [6] to extract information on peptides of interest in a targeted, or peptide-centric, approach from the highly complex MS2 spectra [5,7,8]. This strategy has its roots in the generation of measurement coordinates for use in targeted proteomics [9]. Most commonly this workflow uses an offline fractionation scheme to deeply characterize representative samples using standard DDA data acquisition but requires substantial wet lab and instrument time [10,11]. Methods that attempt to deconvolute the complex DIA MS2 spectra into component MS2 pseudospectra for use in standard database searching approaches are also popular [12,13] but have typically not achieved the same depth of coverage as deep fractionation libraries [14]. Similarly tools that score complex DIA spectra against reference libraries in a spectrum-centric fashion [15], or use a peptide-centric-approach in a library-free strategy [16] have also been proposed. Computationally predicted spectral libraries [17] in DIA analysis have emerged as an alternative to empirically measured libraries, however, a hybrid approach where the predicted libraries are refined using measured data from representative samples has shown the best performance [18–20]. This refinement step serves in part to improve the spectral quality, and in part to reduce search space by discarding undetected peptides. Despite these advances, most published studies still rely on using libraries generated by laborious offline fractionation alone, or as hybrid with the above approaches.

A new approach is to use a gas phase fractionation (GPF) DIA method to improve library coverage without the need for offline fractionation [21]. Searle *et al*, developed a GPF acquisition scheme for library generation for DIA analysis comprised of 4 *m/z* staggered precursor isolation windows arranged so that peptides in the range 400 – 1000 *m/z* are captured [18,22]. This is achieved by six injections of a representative sample covering 100 *m/z* each. This scheme, also integrating empirical refinement of computationally predicted spectra, resulted in data comparable in quality to that derived from a DDA-based fractionation library, showing that GPF can be a reliable method for rapidly developing a sample-specific library. The concepts underpinning GPF-based library generation are applicable to data acquisition using the added dimension of ion mobility separations, such as diaPASEF [23], however, a suitable workflow that coordinates ion mobility separations and GPF is not established.

Trapped ion mobility spectrometry (TIMS) is a practical, sensitive, and selective approach integrating ion mobility spectrometry into quantitative mass spectrometry-based proteomics [24]. In TIMS ions are separated by collisional cross section (CCS) and trapped by counteracting forces of a carrier gas pushing them toward the MS and a ramped electric field that holds them back. Sequential elution of the ions into the MS by reduction in the voltage ramp allow DDA [25] or DIA-based [23] acquisition schemes to benefit from an additional layer of separation and focusing of ions into more concentrated packets. Because CCS and m/z are correlated the mass selective quadrupole can be coordinated in time with the elution from the TIMS cell to maximize efficiency of ion capture in the ddaPASEF or diaPASEF modes of operation.

To determine whether diaPASEF could benefit from GPF based library generation, we implemented a GPF scheme that integrates the ion mobility dimension (IM-GPF) and evaluated its performance using a standard commercial HeLa cell digest sample. We found that our diaPASEF GPF library generation scheme exceeded the performance of libraries generated directly from the diaPASEF data by deconvolution, and had performance approaching that of a deep hybrid library that included ddaPASEF data acquired using basic reversed phase fractionation (BPRF) [23], combined with diaPASEF data and GPF data acquired in this study.

## Results

To determine whether a GPF library generation approach was beneficial for the analysis of diaPASEF data we designed an acquisition scheme integrating narrow m/z windows synchronized with the ion mobility dimension to cover the precursor space used in the ‘standard diaPASEF’ [23] acquisition method (Fig 1). To comprehensively cover the range from 400-1200 m/z we used 5 m/z isolation windows with 1 m/z overlap on both sides requiring 199 quadrupole positions. We allowed 2 quadrupole positions per ion mobility cycle spanning the IM range 0.57-1.47 V/cm^-2^ and split these across 7 acquisition methods with 15 ion mobility cycles (i.e. frames) per method in order to keep the complete duty cycle length similar to that of standard diaPASEF method maintaining sufficient chromatographic peak sampling. To facilitate recalibration procedures across the entire m/z and IM dimensions commonly used in DIA analysis software we distributed the precursor isolation windows evenly across the 7 acquisition methods (and not in adjacent blocks). The coordinates for the 7 GPF methods in the format for direct import into the TimsControl software are provided in the supplementary material.

**Figure 1:**
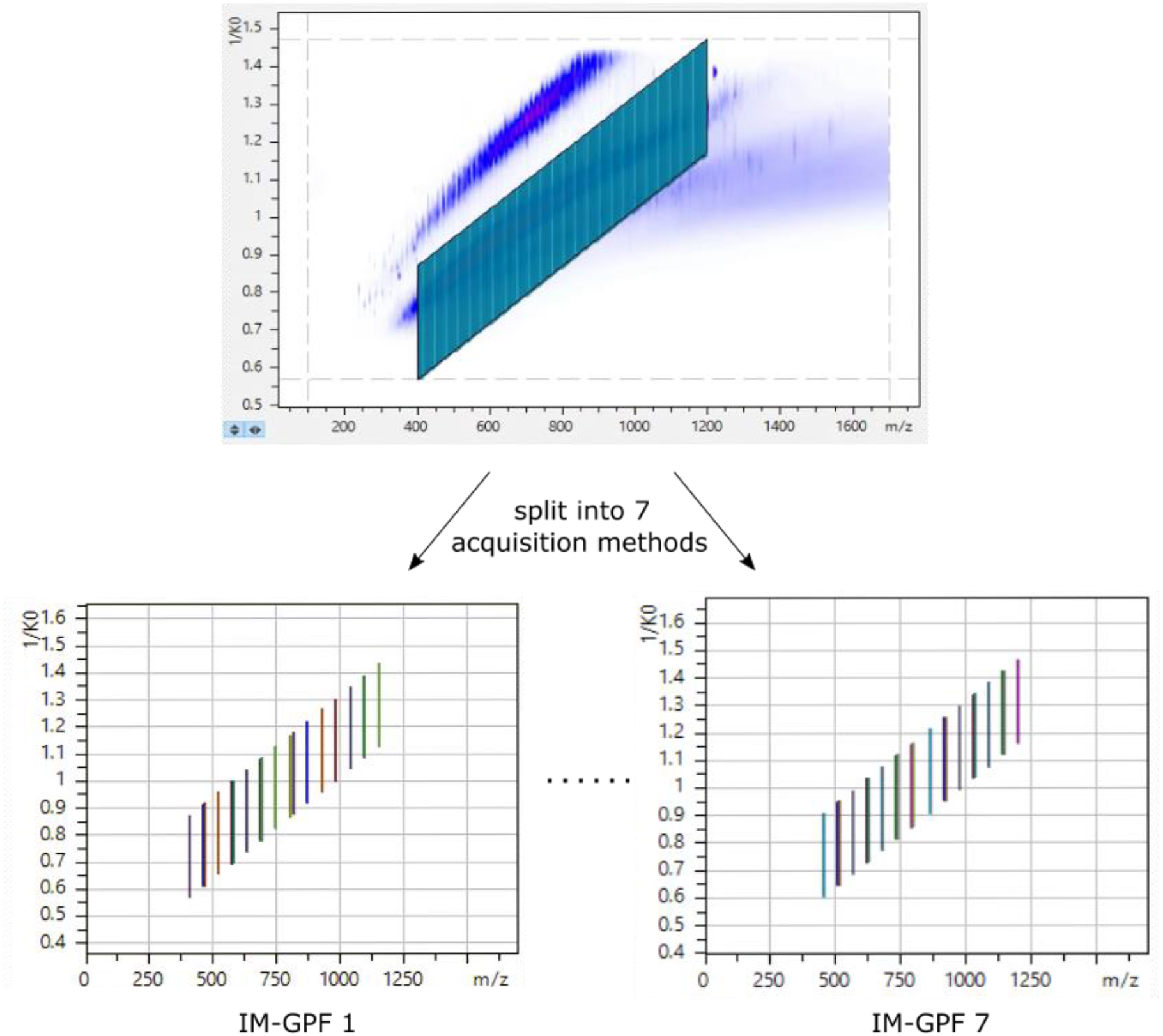
Design of a gas phase fractionation acquisition scheme integrating ion mobility. IM-GPF window scheme 400-1200mz and 0.57-1.47 V/cm^-2^ precursor space using 199 precursor isolation windows. The IM-GPF scheme was split into 7 acquisition methods (representative methods IM-GPF1 and IM-GPF7 are shown).

To evaluate our IM-GPF scheme we constructed 3 spectral libraries from a commercial HeLa digest sample using 3 data types: (i) 3 technical replicate injections using standard diaPASEF acquisition, (ii) the IM-GPF scheme (7 injections), and (iii) a previously published deep HeLa library [23] from basic reversed phase fractionation (24 fractions). The data was processed using the Pulsar search engine integrated in Spectronaut 16 which supports library generation by both direct analysis of DIA data using deconvolution and standard DDA analysis. We refer to these as 1) ‘direct diaPASEF’ – using only the diaPASEF data; 2) ‘IM-GPF’ – using the IM-GPF data combined with the diaPASEF data; and 3) ‘Deep’ – using the BRPF data combined the IM-GPF and diaPASEF data.

Figure 2 shows the composition of these three libraries. As expected, the diaPASEF library was the smallest with 72,447 peptide precursors from 6,524 inferred protein groups. The IM-GPF library contained 82,968 peptide precursors from 6,988 inferred protein groups. This represents an increase of 12.6% and 6.6% over the direct diaPASEF library at the precursor and protein levels respectively. The deep library was substantially larger containing 353,912 peptide precursors from 10,621 inferred protein groups.

**Figure 2:**
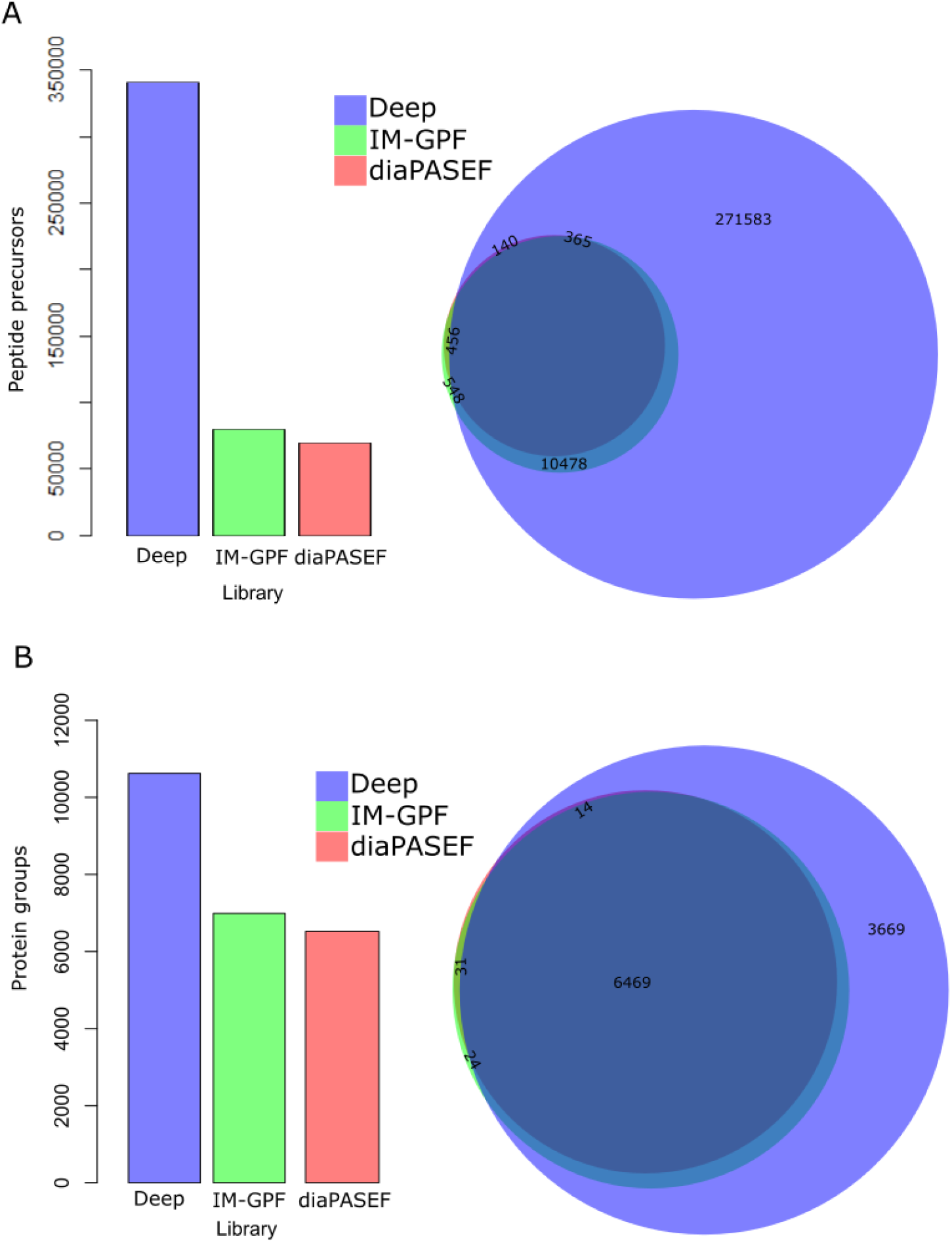
Composition of the direct diaPASEF, IM-GPF, and deep spectral libraries. (A) Peptide precursor identifications in the three libraries (barplots left) and overlap of identified peptide precursors (Venn diagrams right) in the three libraries. (B) Inferred protein group counts from the three libraries (barplots left) and overlap of inferred protein groups from the three libraries.

To assess the difference between the direct diaPASEF, IM-GPF, and deep libraries for the analysis of 3 diaPASEF technical replicates acquired from HeLa cell digests we performed peptide-centric analysis using Spectronaut with each library in turn. Figures 3A and 3B show the number and overlap of the detected peptide precursors and inferred protein groups respectively using each of the three libraries. For the direct diaPASEF library we detected 71,785 peptide precursors and 6,506 inferred proteins in total of which 96.9% and 99.5% could be detected in all three replicates. Using the IM-GFP library we detected 82,170 peptide precursors and 6,981 inferred proteins of which 96.6% and 99.2% were detected in all replicates. This represents an increase of 12.4% peptide precursors and 6.4% inferred proteins detected using the IM-GPF library. Using the deep library we could detect 115,727 peptide precursors and 8,046 inferred proteins in total, however, we note that when considering the number detected in all 3 replicates the IM-GPF library approached that of the deep library detecting with only 8.1% more peptide precursors and 6.3% more inferred proteins identified in all 3 replicates from the deep library. To investigate how the additionally identified peptide precursors were distributed across the abundance range we visualized the peak area for the detected precursors from the 3 libraries as a smoothed histogram (Fig 3C). This analysis showed that while the overall range of peak area values for each of the 3 analyses were similar, the newly identified precursors were primarily found in the lower abundance region as expected. To determine whether quantitative precision was affected by the library used we computed the coefficient of variation (CV) and visualized their distribution as a boxplot (Fig 3D). The CV distributions were comparable although the median CV values were modestly inflated in the IM-GPF and deep library analyses compared to the direct diaPASEF analysis, which is consistent with the recovery of more lower abundance signals.

**Figure 3:**
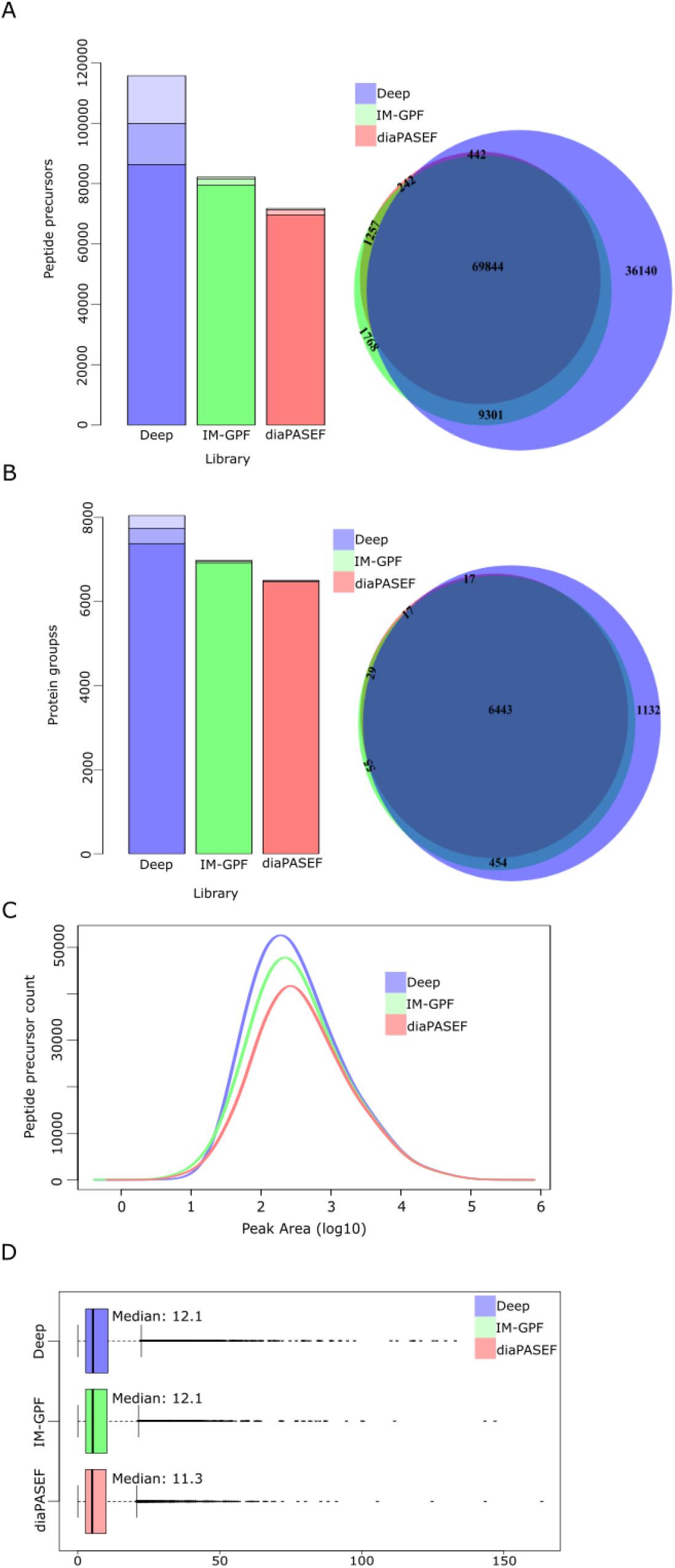
Precursor and protein detection rates in diaPASEF data using different libraries. (A) Peptide precursors detected (barplot left) and overlap (Venn diagram right) in diaPASEF data depending on the library used. The colour shading in ear bar indicates in how many of the 3 replicates the precursor was detected and overlap of detected precursors from each library. (B) Protein groups inferred (barplot left) and overlap (Venn diagram right) in diaPASEF data depending on the library used. (C) Distribution of peak areas for peptide precursors detected with each of the 3 libraries. (D) Distribution of CVs calculated over technical replicates for peptide precursors detected with each of the 3 libraries.

## Discussion

We have developed a gas phase fractionation strategy that integrates trapped ion mobility separations for use in library generation coupled to diaPASEF data analysis. Our scheme has the benefit that no offline separations are required and only 7 additional injections of a representative sample are required similar to the strategy proposed by Searle *et al*. for DIA methods without ion mobility separations [18]. Where potentially heterogenous sample groups are analysed multiple representative sample pools may be included. We show that the performance of the method exceeds that with libraries generated directly from the diaPASEF data in terms of precursor peptide and inferred protein coverage with no effect quantitative precision. While analysis with a deep library based on the more laborious strategy of offline fractionation showed higher total coverage a large fraction of peptide precursors were not detected across all technical replicates. The number of peptide precursors reproducibly detected across technical replicates from the IM-GPF library-based analysis exceeded that of the direct diaPASEF library and approached that of the deep library. As such, we establish IM-GPF as a pragmatic and high-performance solution to generating spectral libraries for use in the diaPASEF data analysis workflow.

## Methods

### diaPASEF data acquisition

Mass spectrometry (MS) proteomics data were acquired on a timsTOF Pro (Bruker) quadrupole time-of-flight mass spectrometer integrating trapped ion mobility separations coupled online via a Captivespray electrospray source (Bruker) to a nanoElute (Bruker) nanoflow liquid chromatography system. All analyses were performed using 200ng HeLa digest (Pierce). Solvent composition at the two channels was 0.1% formic acid in water for channel A and 0.1% formic acid in acetonitrile for channel B. Peptides were separated on an Aurora 2 (Ionopticks) integrated packed emitter column (25cm x 75um ID, 1.6 um C18) using a gradient as follows 2% B, 0 min; 17% B, 60 min; 25 % B, 90 min; 37 % 100 min. The flow rate was 400nl/min and the column was maintained at 50 °C. MS data were acquired in data independent acquisition mode using the ‘standard diaPASEF’ windows scheme (25 m/z width, 4 windows per 100 ms ion mobility scan, 16 ion mobility scans per duty cycle) as described in Meier *et al*^1^.

### IM-GPF data acquisition

The IM-GPF acquisition scheme was designed using the diaPASEF window editor in TimsControl (Bruker). The scheme was designed to cover approximately the same precursor space in the m/z and 1/K0 dimensions as the ‘standard diaPASEF’ [23] acquisition method. We used 5 m/z isolation windows with 1 m/z overlap on both sides requiring 199 quadrupole positions. We allowed 2 quadrupole positions per ion mobility cycle spanning the IM range 0.57-1.47 V/cm^-2^. The windows were exported into text files and then split into 7 acquisition methods where the windows were evenly distributed across 7 acquisition methods with 15 ion mobility cycles (i.e. frames) per method. The 7 schemes were then imported to TimsControl in 7 separate acquisition methods. The coordinates for the 7 GPF methods in the format for direct import to TimsControl software are provided in the supplementary material.

### Data analysis

Library generation was performed using Spectronaut 16 integrating the Pulsar search engine using default settings (automated MS2 demultiplexing; carbamidomethyl cysteine set as fixed modification; oxidized methionine and protein n-terminal acetylation set as variable modification; fully specific Trypsin/P digestion; 2 missed cleavages; peptide length 7-52; mass tolerances set to dynamic, precursor and protein Qvalue set to 0.01). DIA data analysis was performed using Spectronaut 16 using default settings (mass tolerances, IM extraction window, and RT extraction window set to dynamic; decoy method mutated; precursor and protein Qvalue set to 0.01; data filtering set to Qvalue; imputation switched off). Spectronaut results at peptide precursor and protein group level were exported and analysed with R using the Biovenn package and base functions boxplot and density.

## Supporting information

Window Coordinates for IM GPF

## Acknowledgements

JP was supported by a Co-operative Awards in Science and Technology studentship funded by the Northern Ireland Department for the Economy and Bruker Daltonik GMBH. We thank Stephanie Kaspar Schoenfeld and Nagarjuna Nagaraj (Bruker Daltonics) for helpful discussions.

## Data availability

The mass spectrometry proteomics data have been deposited to the ProteomeXchange Consortium via the PRIDE [26] partner repository with the dataset identifier PXD034640. Raw data for the previously published BRPF HeLa library [23] are available with the dataset identifier PXD017703 (see HeLa_200ng_Library_RAW.zip).

